# Cross-linking mass spectrometry for structure analysis of the intrinsically disordered Tau and phosphorylated Tau protein

**DOI:** 10.1101/2025.09.02.673744

**Authors:** Cristian Arsene, Alexander Gates, Anne-Katrin Römmert, André Märtens, Valentina Faustinelli, Luise Luckau, Gavin O’Connor

**Affiliations:** Physikalisch-Technische Bundesanstalt (PTB), Braunschweig und Berlin, D-38116 Braunschweig, Germany; School of Data Science, University of Virginia, Charlottesville, VA 22904, USA; National Measurement Laboratory, LGC, The Priestley Building, 10 Priestley Road, Guildford, GU2 7XY, UK

## Abstract

We present a novel method for analyzing the folding of intrinsically disordered proteins (IDPs), such as Tau and phosphorylated Tau (pTau), in solution. Using cross-linking mass spectrometry combined with a new downstream analysis framework, we construct weighted interaction networks from cross-link–derived residue pairs without relying on predefined secondary structure assumptions. Structural differences between protein conformations are quantified by comparing the organization of loop structures within their cross-link networks. Validation with bovine serum albumin (BSA) in native and denatured states shows that at least 500 cross-links—requiring 5–10 replicate measurements—are needed for reliable detection of structural divergence. Leave-one-out analysis confirms that structural transitions are global, highlighting the importance of comprehensive cross-link datasets. The coverage of unique cross-links was evaluated using accumulation curves from randomized permutations. Saturation levels were found to be 9.7%, 5.0%, and 6.2% of the total 528 and 10,731 possible cross-links after 30, 84, and 62 technical replicates, respectively, for myoglobin, native BSA, and denatured BSA. For Tau and pTau, coverage reached 10.8% and 5.5% of the upper limit (8,256). Finally, applying our structural analysis to Tau and pTau during arachidonic acid–induced aggregation revealed distinct patterns of structural evolution between the two proteins.

## Introduction

Particularly for intrinsically disordered proteins, where structures cannot be elucidated using x-ray chrystallography or cryo-EM, alternative methods still enable the analysis of the three-dimensional protein structure at least at a moderate spacial resolution. As one of those methods, cross-linking mass spectrometry (XL-MS) has evolved into an important tool for the analysis of protein structures in solution.^1^ Intrinsically disordered proteins tend to aggregate under pathological conditions and are involved in the development of neurodegenerative diseases. Neurofibrillary tangles of the aggregated Tau protein in brain cells are a hallmark of Alzheimer disease and other tauopathies. ^2^ In this context, hyperphosphorylation of Tau is discussed as a trigger for misfolding of the protein and the formation of oligomers, protofibrills and finally insoluble neurofibrillary tangles. For biomarker proteins of neurodegenerative diseases, including Alzheimer’s disease, it is particularly important to understand the reason for changes in three-dimensional structure and their impact on the tendency of such proteins to aggregate and to form cytotoxic intracellular neurofibrillary tangles. Here, we propose a methodology for the identification of discrepancies in the higher-order structure of proteins. To illustrate this approach, we utilise an example of the phosphorylation status of a recombinant Tau protein in solution, distinguishing between phosphorylated (pTau) and non-phosphorylated states (Tau). XL-MS was used for structure analysis, providing moderate but otherwise inaccessible information about distance constraints and the three-dimensional structure of proteins in solution. Myoglobin and bovine serum albumin (BSA), two proteins of different sizes, were used to optimize the XL-MS protocol. BSA was chosen as a model for the development of the structure analysis method. Defined conditions were adjusted to induce changes of the three-dimensional structure of BSA. Covalently binding cross-linkers were used to “freeze” the induced structures and bottom-up proteomics was adapted for the analysis of cross-linked peptides. Maps of cross-linked amino acid positions in the protein sequence were generated as intra-molecular loop-networks for BSA in its native and denatured form and each loop-network was submitted to unsupervised cluster analysis. Differences in native and denatured structures were detected by pairwise comparison of their cluster analysis. Apart from the detection of structure changes at the global level, it is possible to pinpoint sequence stretches, of different higher-order structures, by plotting the element-wise agreement in the pair-wise clustering comparison. Performance measures for the clustering comparison were calculated using a bottom-up approach and recently developed statistical tools, included into the Python package CluSim. ^3,4^ This measurement principle was applied to the phosphorylated/non-phosphorylated Tau proteins in solution. The changing folding states of Tau and phosphorylated Tau were investigated by analysing aliquots of protein solutions at different time points on the time line of in-vitro Tau-aggregation. The loop-network developed from cross-linking data for the protein structure at the beginning of the kinetic series was used as reference and pair-wise compared to the loop-network of cross-linking data obtained at different time points of aggregation.

## EXPERIMENTAL SECTION

### Protein materials

#### Native BSA

BSA was purchased from Sigma-Aldrich, St. Louis, USA, cat. no.: 05470. A solution of 10 *μ*M BSA in 1 mL of 50 mM 4-(2-Hydroxyethyl)-1-piperazine ethanesulfonic acid (HEPES)-buffer (pH7.5) was prepared and filtered by centrifugation at 7000 *xg* through a 2 mL-Amicon Ultra centrifugal filter (10 kDa MWCO). The residue on the Amicon filter was washed 5 times with 50 mM HEPES-buffer, and finally dissolved in 1 mL of 50 mM HEPES-buffer and transferred to cross-linking.

#### Denatured BSA

For the structure analysis of denatured BSA, an additional solution of 10 *μ*M BSA was prepared in HEPES-buffer. For reduction of disulfide bridges, an aliquot of 10 *μ*L of an aqueous 34 mM dithiothreitol (DTT) solution was added and the protein solution was incubated for 1 hour at 60 ° C. For alkylation, an aliquot of 10 *μ*L of an aqueous 34 mM 2-iodoacetamide (IAA) solution was added and the protein solution was incubated for 30 min at room temperature. This solution was filtered through a 2 mL-Amicon Ultra centrifugal filter and washed as the native-BSA solution.

#### Myoglobin

Myoglobin from horse skeletal muscle was obtained from SIGMA-Aldrich, St. Louis, USA, cat. no.: 70025). A solution in 50 mM HEPES-buffer was prepared as described for denatured BSA.

#### Tau and phosphorylated Tau (pTau)

Recombinant Tau and pTau (GSK-3*β*-phosphorylated) were from Acro Biosystems, Basel, Switzerland. Both materials were obtained as solutions of 0.5 mg/g in 50 mM Tris, 150 mM NaCl, 1 mM DTT, 1 mM EDTA, pH 7.5. The phosphorylation pattern of pTau was analysed using a standard bottom-up proteomics protocol, as recently reported. ^5^ Phosphorylation was used as variable modification for database search. Intact protein mass spectrometry was used to investigate the distribution of the number of phosphorylations in pTau, as previously described. ^6^ The material contained unphosphorylated Tau beside uniformly triphosphorylated Tau. Intact protein mass spectrometry revealed a deconvoluted mass of 45718.7 and 45953.4 Da, for Tau and pTau respectively. Expected masses were: 45716.7 (Tau) and 45958.7 Da (pTau). The pTau material was further purified using strong-anion exchange chromatography on a MONO-Q 4.6/100 PE column. Gradient elution was applied within 42 min between 0 and 20% NaCl in a 20 mM Tris-buffer. The flow rate was 1mL/min. Fractions containing pTau were collected. The stock solutions of Tau and the purified pTau were filtered and washed in the same way as the BSA solution. Aliquots of Tau or pTau solution containing *∼* 1 nmol of the protein were submitted to aggregation reaction and cross-linking.

### Aggregation of Tau and pTau

For aggregation, 2 mL of a solution of 4 *μ*M Tau or pTau were prepared in a buffer, containing 50 mM HEPES and 100 mM NaCl at pH 7.5. A sample (500 *μ*L) was taken and the aggregation was started by addition of 131.3 *μ*L of 2 mM arachidonic acid (ARA) in ethanol to the remaining Tau or pTau solution. Additional samples (500 *μ*L) were collected at 30, 90 and 150 sec following the initiation of the aggregation. From each sample, an aliquot of 10 *μ*L was taken for the Tau aggregate ELISA. Each remaining sample was diluted in 500 *μ*L of the 50 mM HEPES buffer, containing 100 mM NaCl and submitted to the cross-linking reaction.

### Cross-linking

The cross-linking reagent disuccinimidyl dibutyric urea (DSBU) was obtained from ThermoFisher Scientific, Darmstadt, Germany, cat. no.: A35459. A stock solution of 25 mM DSBU in 94 *μ*L of dried DMSO was prepared. To each filtered protein solution or aliquot from aggregation solution a 20 *μ*L-aliquot of the cross-linker stock solution was added and gently shaked for 1 hour at room temperature. The reaction was stopped by addition of 20 *μ*L of 1M ammonium hydrogen carbonate solution. The solution was confined under vacuum at room temperature and submitted to proteolysis.

### Proteolysis

To the product of the cross-linking reaction 25 *μ*L of denaturation buffer were added, containing 8M urea in 400 mM ammonium hydrogen carbonate. The solution was denatured for 5 min in an ultrasonic bath. For reduction of disulfide bridges, an aliquot of 8 *μ*L of an aqueous 20 mM DTT-solution was added and the reaction solution was incubated for 30 min at 60 ° C. For alkylation, an aliquot of 10 *μ*L of an aqueous 60 mM IAA-solution was added and the reaction solution was incubated for 20 min at room temperature. The excess of IAA was quentched by addition of 36 *μ*L of a 20 mM DTT-solution. Pure water (160 *μ*L) was added to lower the urea concentration to 1 M and the proteolysis was started by addition of 13 *μ*L of a solution containing trypsin at 1*μ*g/*μ*L. An additional aliquot of 13 *μ*L of the trypsin solution was added after 24 hours of incubation at 37 ° C and the proteolysis was continued over night. The product was desalted on solid phase cartridges following a standard protocol using two 5 mM ammonium acetate buffers: an aqueous buffer for the wash step and a solution of 80% acetonitrile for the elution of peptides. The eluate was confined under vacuum and the residue of 60 *μ*L was submitted to LC-MS/MS analysis.

### Liquid chromatography-mass spectrometry

An UltiMate 3000 RSLCnano HPLC system (Thermo Fisher Scientific) coupled to a timsTOF Pro mass spectrometer (Bruker Daltonics) was used for the analysis of the proteolysed sample of cross-linked protein. Aliquots of 2 *μ*L were analysed. Peptides were trapped on a pre-column (Acclaim PepMap C18, 5 *μ*m, 0.3×5mm) and then separated on a Bruker Fifteen nanoFlow column (25cm x 150 *μ*m, C18, 1.9 *μ*m, 120 Å) using a water-acetonitrile gradient from 2 to 17% B in 55 min, from 17% to 25% B in 30 min then from 25% to 37% B in 10 min and from 37% to 90% B in 10 min (with solvent A: water, 0.1 vol.-% formic acid and B: acetonitrile, 0.1 vol.-% formic acid) at 50 °C. The flow rate was 700 nl/min. The timsTOF Pro mass spectrometer was equipped with a CaptiveSpray ion source. The mass spectrometer was run using the DDA-PASEF-standard-1.1 sec-cycletime method, as provided by Bruker with some modifications. Briefly, the settings were: 14 PASEF MS/MS scans per acquisition cycle with a trapped ion mobility accumulation and elution time of 166 ms. The charge state minimum and maximum for precursor ions was set to 3 and 8, respectively. Spectra were acquired in a *m/z* range of 100 to 1700 and in an (inverse) ion mobility range (1*/K*_0_) of 0.60 to 1.60 Vs/cm^2^. The collision energy was set up as a linear function of ion mobility starting from 20 eV for 1*/K*_0_ of 0.6 to 95 eV for 1*/K*_0_ of 1.6.

### Database search for cross-links

MeroX 2.0, ^8^ Version 2.0.1.4 was used for identification of cross-linked peptides. Databases for *BSA, horse myoglobin* and *Tau* were obtained as FASTA files (uniprot.org, P02769, P68082 and P10636-8, respectively for BSA, horse myoglobin and microtubule-associated human protein tau, accessed: 18. Apr. 2023). The following settings were applied for data analysis: carbamidomethylation of cysteine as fixed modification, methionine oxidation and phosphorylation of amino acids serine, threonine or tyrosine as variable modifications. Proteolytic cleavage sites were R and K and DSBU was set as cross-linking reagent for the reaction at side chains of amino acids lysine, serine, threonine and tyrosine and at the N-terminus of the protein. The mass tolerance for the monoisotopic mass of precursor ions and fragment ions was 15 and 25 ppm, respectively. Only *b−* and *y−* fragment ions were considered. Mass correction was enabled for precursor ions. The minimum score for cross-linked peptides was set to 20. The false discovery rate (FDR) was 1%. Lists of identified cross-linked peptides including the cross-linked amino acid positions within the protein sequence were used for downstream data analysis.

### Cross-link detection saturation analysis across replicated measurements

To assess cross-linking site coverage across measurements, replicates of datasets were randomly permuted and processed to extract non-homeotypic cross-links. After filtering, site-pairs were converted to unique combinations and counted cumulatively over increasing numbers of technical replicates. This was repeated across 84 cycles (for native BSA), and the mean number of detected unique cross-links was computed. Parallel simulations using randomized site-pair permutations were performed over 199 cycles (for native BSA) to estimate stochastic accumulation of cross-links under uniform random sampling. Both empirical and simulated distributions were visualized using Matplotlib with incremental sampling plotted as red and black scatterlines, respectively.

### Parsing of cross-links

From the list of identified non-homeotypic cross-linked peptides the pairwise cross-linked lysine, serine, threonine and tyrosine positions within the protein sequence were parsed and merged into a list of cross-linked sequence sites.

### Loop-based clustering similarity of protein structures

Using crosslink-derived sequence site pairs and the inherent connectivity of amino acid residues, weighted interaction networks were constructed using the Python package NetworkX. ^9^ Specifically, proteins are represented as networks in which nodes correspond to amino acid residues and edges capture two distinct types of interactions: (1) physical neighbor relationships based on the primary sequence, and (2) cross-linking interactions identified experimentally. The resulting graph was analyzed to detect loop structures, i.e. closed paths formed by cross-links that bridge distant regions of the protein. This analysis provided insight into folding patterns and structural constraints within the disordered ensemble. Each loop is treated as an overlapping cluster, allowing for residues on loop endpoints to have multiple memberships in adjacent loops. In order to compare loop-based clustering across conditions (e.g., native vs. denatured BSA or Tau vs. pTau), an element-centric similarity (*α* = 0.9) was applied using the CluSim Python package ^3,4^ Bootstrap resampling (n = 10) across increasing data subsets was performed to assess classification consistency. Resulting similarity metrics were visualized as error-bounded line plots to reveal intra- and inter-sample variability.

### 3D Visualization of node-specific similarity in protein networks

Normalized element-centric similarity scores between loop-clustered native and denatured BSA or Tau and pTau networks were visualized in 3D space. A weighted multigraph, derived from cross-linking data and stored in GML format, was processed using NetworkX. Node positions were computed via spring layout embedding (dim=3, iterations=1000), with edge lengths modulated to reflect cross-link constraints. To ensure reproducibility of node placement, a fixed random seed (seed=27061971) was applied to the layout algorithm. Element-specific similarity scores (*α* = 0.9) from CluSim were mapped to node color using a plasma colormap and visualized using Plotly. Short and long edges were rendered separately to distinguish structural patterns, and every 10th node was annotated for orientation.

### Leave-one-out analysis of cross-link influence

To evaluate the structural impact of individual cross-links unique to native BSA (Tau), a leave-one-out approach was applied. Each cross-link was removed from the native BSA (Tau) network one at a time, and the resulting network was reanalyzed to identify changes in loop topology. The modified network was compared to both the original native BSA (Tau) and denatured BSA (pTau) networks using element-wise clustering similarity scores (CluSim, *α* = 0.9).

## Data availibility

The mass spectrometry proteomics data have been deposited to the ProteomeXchange Consortium via the PRIDE partner repository with the dataset identifier PXD067639.

## RESULTS AND DISCUSSION

### Identification of cross-links and distance-constraints

Building on an established XL-MS protocol, ^10^ liquid chromatographic conditions were further refined to enhance the reliable detection of cross-linked peptides following protein digestion across a larger number of technical replicates. The workflow as shown in Fig. 1 was validated by analysis of horse myoglobin and bovine serum albumin (BSA), two model proteins with different amounts of reaction sites for cross-linking. The maximum number of cross-links between lysines, serines, threonines and tyrosines is 528 and 10731 in myoglobin and albumin, respectively. However, only a small fraction of 9.7% and 5.0% of the total number of cross-links was identified across 30 and 84 technical replicates in myoglobin and in native BSA, respectively (Fig. 2A,B). To overcome the limited detection of cross-linked peptides in single MS runs, observed in our optimised protocol for myoglobin and BSA and consistent with Belsom et al. ^11^, each sample was analyzed in at least 30 technical replicates. This approach balanced analysis time with data completeness, allowing software-assisted ^8^ identification of the majority of cross-linked amino acid site pairs within the protein sequence (Fig. 2A,B).

**Figure 1:**
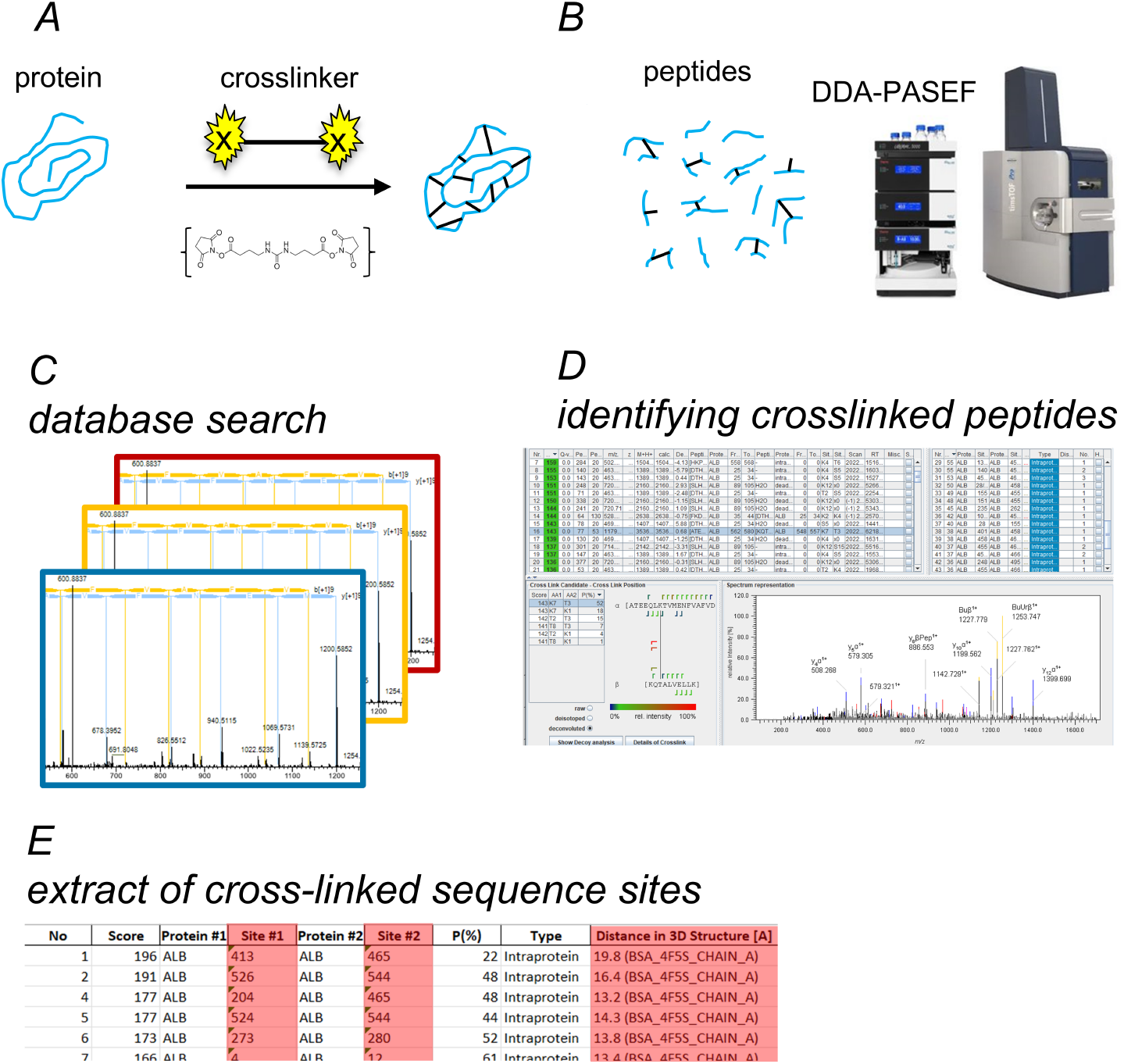
Workflow for cross-linking MS analysis. (A) The pure protein is cross-linked by reaction with DBSU. (B) The cross-linked protein is digested and tryptic peptides are subjected to data dependent analysis using the standard DDA-PASEF method included in the instrument-control software of the TIMS-ToF mass spectrometer. (C) Cross-linked peptides are analysed by data base search for precursor- and fragment ions using the MEROX software. (D) Cross-linked petides are identified based on default statistical criteria. (E) Cross-linked sequence sites are filtered from the result list of data analysis for exclusion of homeotypic interactions and decoy entries. If available, data from X-ray crystallography can be used to compare cross-linked sequence sites, allowing distance constraints to be extracted from the MEROX results list

**Figure 2:**
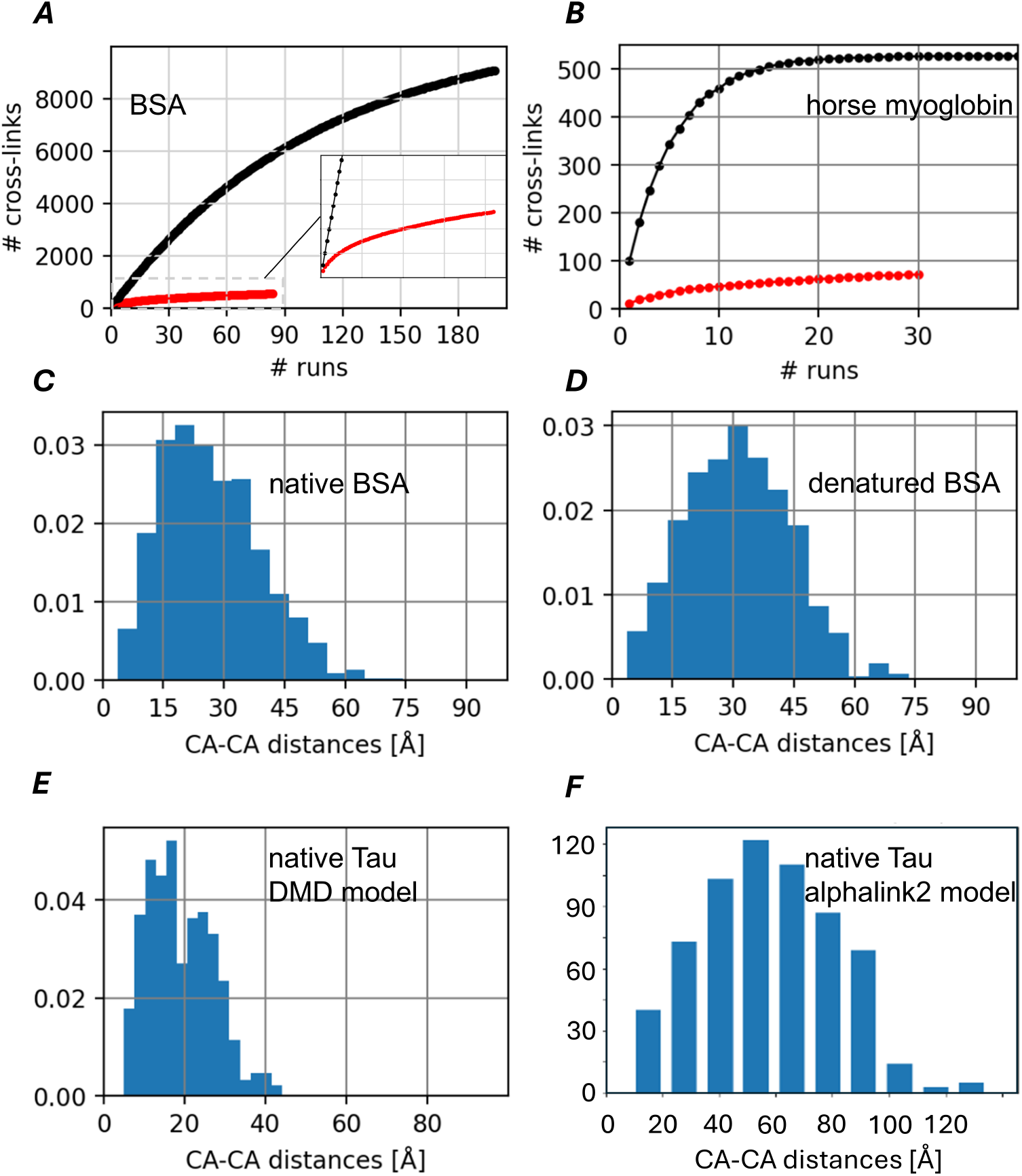
Accumulation of unique cross-links across technical replicates and the distribution of distance constraints. (A, B) The number of identified unique cross-links (red line) was accumulated across randomized permutations of technical replicates of the same cross-linked and digested protein. The black line shows the expected accumulation of unique cross-links based on a random sampling model, assuming 100 randomly selected links per cycle from a fixed pool of possible cross-links. (C-F) Distributions of distance constraints were derived by comparing the identified cross-linked sequence sites from native and denatured BSA and from native Tau to their corresponding Euclidean distances between respective C-alpha atoms in the X-ray crystal structure of BSA or in the Tau conformer, either predicted by discrete molecular dinamics (DMD) simulation or by alphalink2.

Duplicates were not eliminated, since the amount of duplicates consists chemical information about accessibility of reaction sites for cross-linking and therefore structural information. As a test for the analysis of structural changes of the protein folding, BSA was analysed in its native form and after protein denaturation. In denatured BSA, as with native BSA, only 6.2% of the total cross-links were detected across 62 technical replicates (data not shown). The identified cross-links in both native and denatured BSA were aligned with the distances between cross-linked sequence sites derived from the X-ray crystal structure of native BSA. ^12^ The distributions of these distance constraints are shown in Fig. 2C, D. Compared to native BSA, the distribution for denatured BSA is shifted toward longer distances between cross-linked sites. This indicates a more flexible and extended conformation of the denatured protein.

### Set-up of loop-networks as models for protein structures

Attemps were made to obtain higher-order structural insights in protein folding by integrating XL-MS data with structural predictions resulting in the development of alphalink2. ^13^ However, the analysis of the intrinsically disordered Tau protein using alphalink2 predominantly yields, as already expected from alphafold prediction, ^14–16^ a random coil conformation, with only a few short helical segments (data not shown). Discrete molecular dynamics (DMD) simulations of the Tau conformer was also combined with cross-linking data resulting in a predicted globular protein model. ^17^ In comparison to the alphalink2 model which we obtained by allignment of our own cross-linking data with the predicted protein conformer, the globular Tau model displays only limited regions of unstructured folding. Its predominantly globular structure and the abundance of secondary structural elements do not adequately account for the intrinsic flexibility of Tau, which is critical for its dynamic association and dissociation with neuronal microtubules under physiological conditions. The resulting distribution of distance constraints for the alphalink structure model was broad, consistent with the behavior expected from a highly flexible protein conformation. For comparison we alligned our cross-linking data also with the globular Tau model predicted by DMD simulation, resulting in a narrow distribution of short distance constraints (Fig. 2E,F).

A resolution to this inconclusive scenario may be achieved by obviating the need for assumptions regarding secondary structure, which are often difficult—or even impossible — to make for intrinsically disordered proteins. Here, we recast the comparison of protein structures in terms of loops induced by cross-links within a topological network model. This approach is based on the premise that cross-links between lysine, serine, threonine, or tyrosine residues introduce bends in the protein’s primary structure, thereby forming looped configurations.

Specifically, we assessed the structural similarity of BSA in its native and denatured forms by performing pairwise comparisons of clusterings derived from the corresponding loop networks. The CluSim package ^18^ was used to compute element-wise similarity scores, providing quantitative measures of similarity at individual sequence positions, where each position was treated as a cluster element. (Fig. 3A,B).

**Figure 3:**
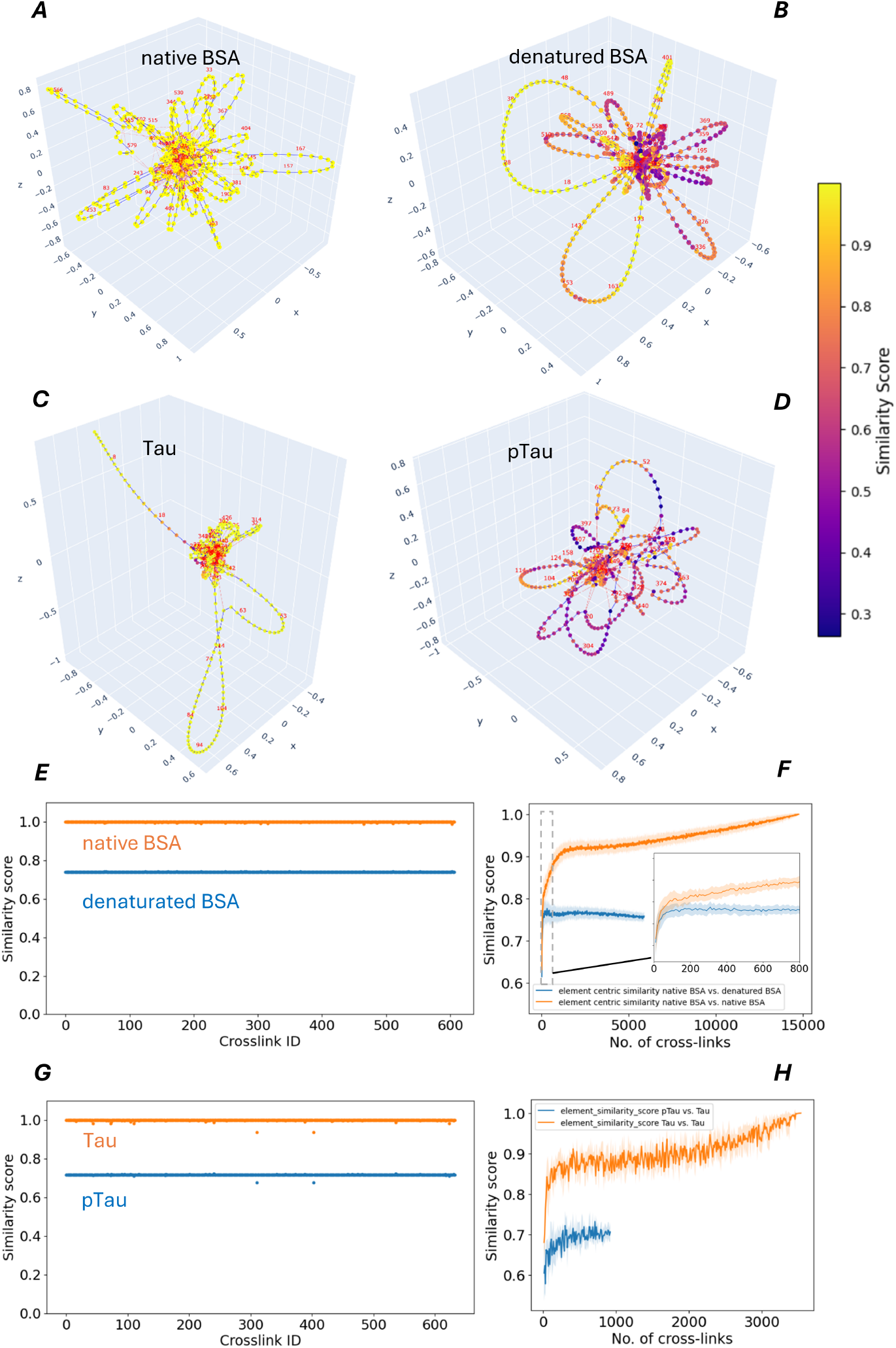
Cross-link networks of BSA, Tau, and pTau conformers, and the element-centric similarity of their loop clusterings. (A − D) Clustering similarity based network visualizations of cross-link networks for BSA, Tau, and pTau conformers, with pairwise comparisons of native vs. denatured BSA using native BSA as the reference (A,B) and Tau vs. pTau using Tau as the reference (C,D). Structural differences are highlighted via element-wise similarity scores between loop clusterings. Red edges denote cross-links (edge length= 8), and blue edges represent sequential (primary sequence) connections (edge length = 1). (E,G) Distributions of similarity scores across sequence positions using a leave-one-out model, where for BSA each native-specific cross-link and for Tau each Tau-specific cross-link is removed to assess its influence on clustering similarity to both protein pairs (native and denatured BSA, or Tau and pTau). (F,H) Element-centric similarity of loop clusterings derived from cross-link networks of native vs. denatured BSA (F) and Tau vs. pTau (H). Similarity scores are shown as a function of sample size, averaged over randomized subsamples of cross-links; shaded areas represent standard deviation.

To determine whether individual cross-links account for the primary differences between the two protein conformers, we first identified the extra cross-links present exclusively in native BSA. These were then removed one at a time using a leave-one-out approach, after which the resulting loop clustering was compared both to the original network and to that of denatured BSA to calculate element-wise similarity at each sequence position (Fig. 3E). Similarily, cross-link data obtained for Tau and pTau were used to construct the respective topological networks and extract their loop clustering structure, which were analyzed in the same manner as the BSA networks (Fig. 3C,D). Using the leave-one-out approach developed for the BSA networks, we evaluated the influence of individual cross-links on structural similarity between Tau and pTau. Notably, removal of two Tau-only cross-links reduced the similarity of the Tau loop network to its original form and, unexpectedly, also decreased its similarity to the pTau loop network. Given that these cross-links were considered key differentiators between the two networks, we had anticipated that their removal would increase similarity to pTau, not decrease it. Plots of element-similarity scores for both BSA and Tau (Fig. 3E,G) indicate that such changes affect the protein s overall structure, underscoring the need for a holistic analysis of cross-links to accurately capture structural features. The limited subset of cross-links detectable in a single measurement is insufticient for this purpose. In Tau and pTau, only 10.8% and 5.5% of the maximum possible cross-links (8,256) were detected across 100 and 120 technical replicates, respectively (data not shown).

### Sample size and associated error for pair-wise comparison of clusterings

It can be emphasized that the size of cross-link subsamples has an influence on the pair-wise analysis of loop-network clusterings. To investigate the lower limit of sample size for the detection of divergence between the cross-link networks for both BSA conformers or for Tau and pTau, randomized cross-link subsamples of increasing size were pair-wise analysed, based on clustering of the corresponding loop-networks (Fig. 3F,H). For BSA, the minimum sample size of cross-links, which is necessary to reliably detect structural divergence between both conformers is 500 with duplicates included. Assuming the detection of 50-100 cross-links per technical replicate, at least 5-10 repetitions would be necessary.

### Structure changings of Tau and pTau in the run-up to aggregation

The aggregation of Tau and pTau was induced by addition of arachidonic acid (ARA) as previously described. ^19^ For monitoring the time course of aggregation, a Tau aggregate ELISA was used, which sensitively detects oligomers and aggregates ^7^(experimental section). The epitope of the assay antibody is located in the Tau amino acid sequence region of 428 to 437. While the aggregation of Tau is saturated after 2 hours reaction time, the aggregation of pTau is inhibited (Fig. 4A). There is no direct interference of the assay antibody with the phosphorylation sites, which may explain the very low ELISA results for pTau aggregates, since our pTau material is uniformly threetimes phosphorylated at T169, S199 and T217, while the epitope of the assay-antibody is far away from this region. At the beginning of the aggregation, aliquots were taken and immediately cross-linked for structural analysis. The networks and element-centric similarities of tau cross-linked at different time points during aggregation, and compared pairwise with tau cross-linked before the start of aggregation, are shown in Fig. 4B-E. The structure of the Tau conformer significantly changed right at the beginning of the aggregation and its structural divergence remained detectable at the following time points (90 and 150s). The cross-linking reaction was optimized for the formation of intra-molecular cross-links by chosing a low protein concentration. However, during aggregation, the formation of intermolecular cross-links within oligomers of Tau cannot be omitted. Therefore, aliquots for cross-linking were taken only at the beginning of the aggregation within 2-3 min of the reaction time when oligomers and aggregates are not yet formed.

**Figure 4:**
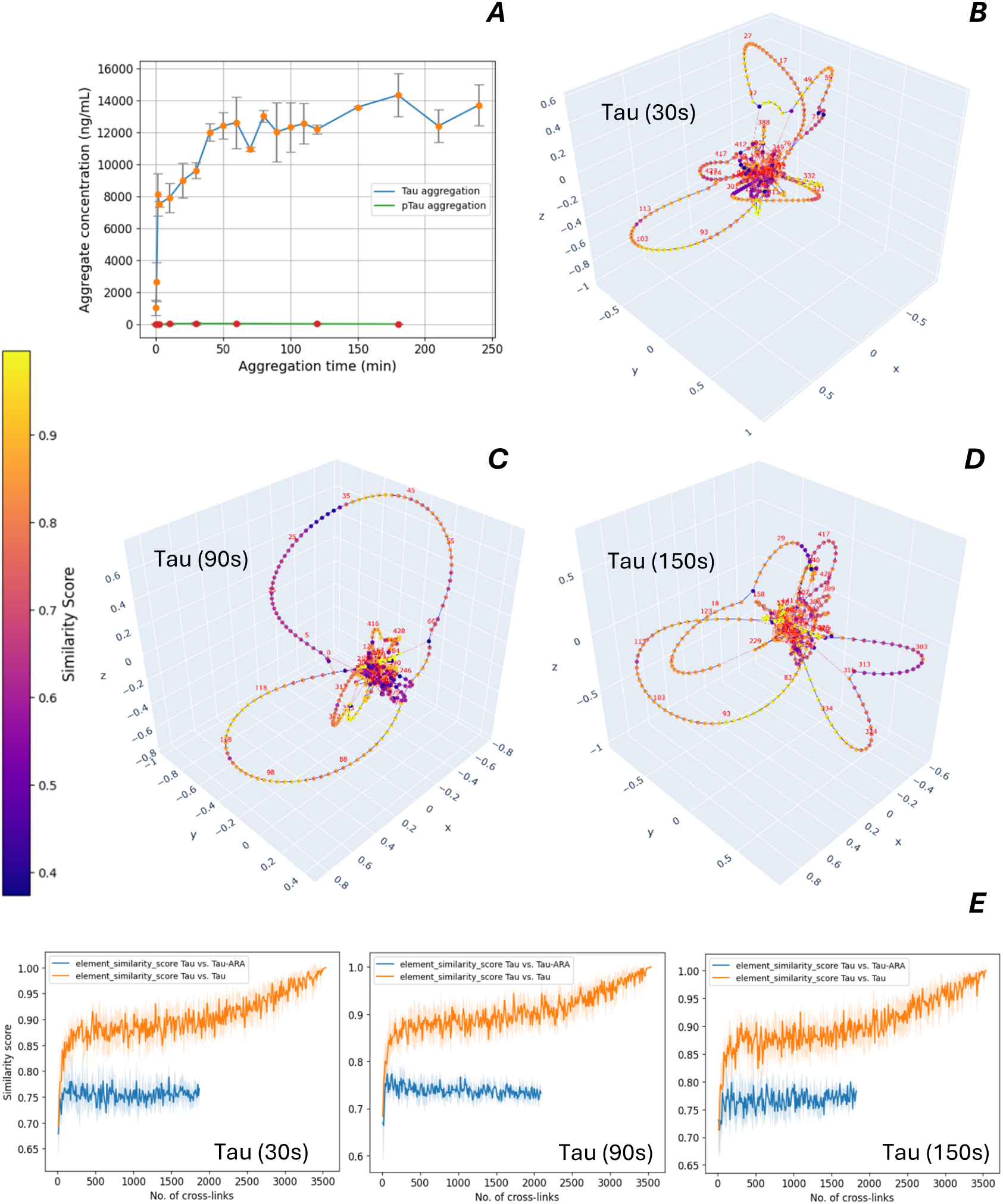
Aggregation and structure analysis of Tau conformers. (A) Time course of Tau and pTau aggregation, induced by arachidonic acid. An ELISA was used to quantify the tau aggregates. Mean values are shown with error bars corresponding to the standard deviation (n=2). (B–D) Cross-link network visualizations for Tau at different time points during aggregation. (E) Similarity scores for the loop clusterings as a function of sample size, comparing the cross-link network of each Tau conformer (aggregated for different durations) to the reference Tau network obtained prior to aggregation with arachidonic acid. Scores are averaged over randomized subsamples of cross-links; shaded areas represent standard deviation.

The same analysis method was applied to pTau and its aggregation kinetics. If compared to the aggregation of Tau, only small changes of the structure were detected during the preliminary phase before aggregation (Fig 5A-F). In contrast to our triphosphorylated pTau material, the hyperphosphorylated Tau, which was aggregated under similar conditions (75 μM ARA, 2 μM Tau or pTau) in a previuos study, ^20^ showed very similar aggregation kinetics as unphosphorylated Tau. However, the electrostatic environment of at least 12 phosphorylated sites across the seqence of the hyperphosphorylated Tau influences the higher-order protein structure and its flexibility most probably in a different way, resulting even in the opposite aggregation behavior, if compared to our triphosporylated Tau material.

**Figure 5:**
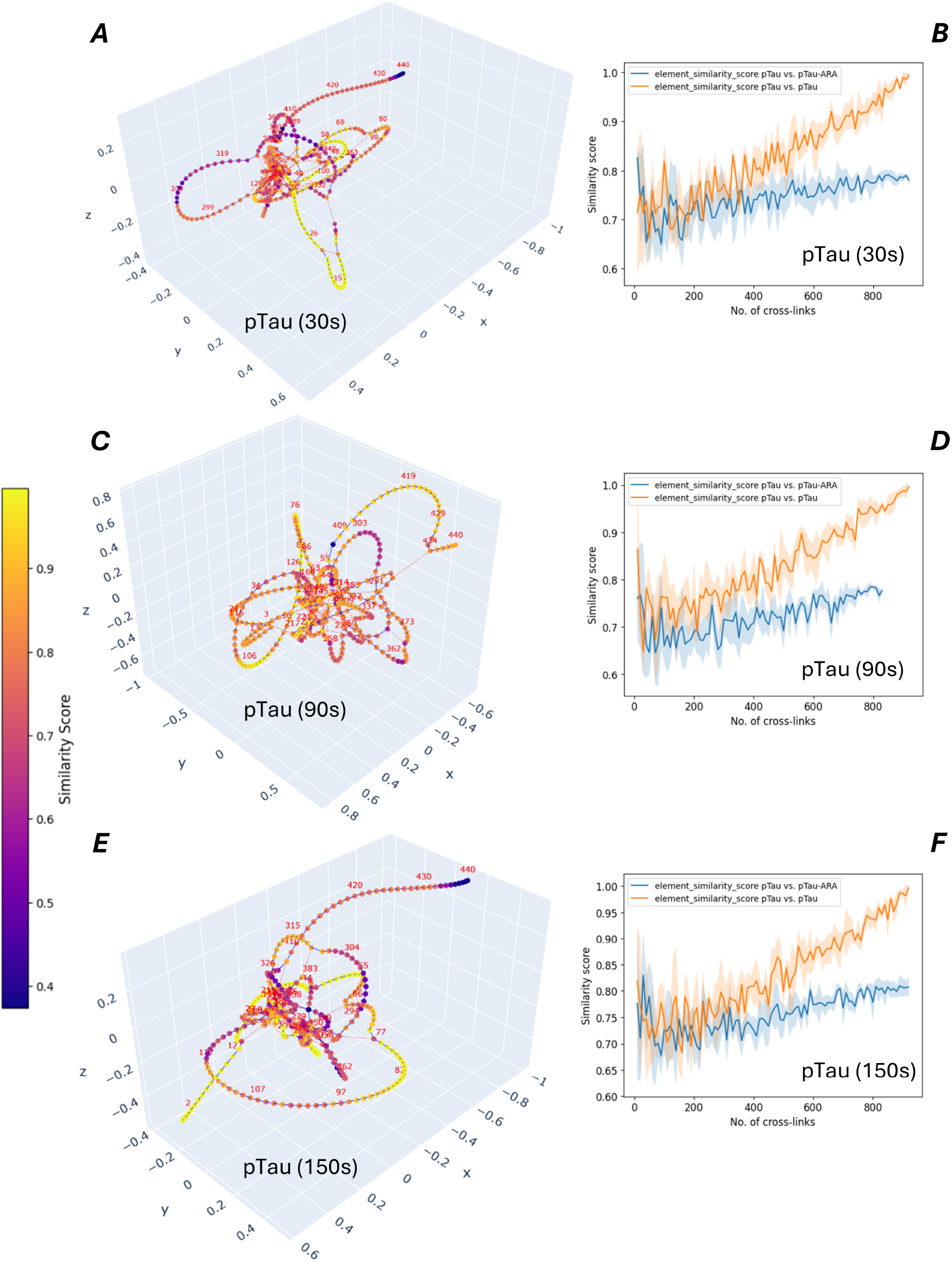
Aggregation and structure analysis of pTau conformers. (A,C,E) Cluster-based network visualizations of cross-linked pTau at different time points of aggregation. (B,D,F) Similarity scores are shown across sample sizes for the comparison of the network of each pTau conformer, cross-linked at different time points of aggregation, and the reference network for pTau, cross-linked at the beginning before starting the aggregation with arachidonic acid. Similarity scores were averaged over randomized subsamples of cross-links. Shaded areas indicate standard deviation.s

## CONCLUSIONS

The new cross-linking analysis method enables the sensitive detection and relative quantification of structural divergence between protein conformers under different conditions and over time. This opens the possibility to further investigate the influence of post-translational modification (PTM) patterns e.g. the differential phosphorylation of Tau on the relationship between structure and aggregation. The XL-MS based structure analysis may also be paired with activity assays for quality control of therapeutic proteins by comparing sample lots to a reference batch with established activity.

## Acknowledgements

This research has received funding from the European Partnership on Metrology, cofinanced from the European Union’s Horizon Europe Research and Innovation Programme, and by the Participating States (European Partnership on Metrology, 10.13039/100019599, 22HLT07 NEuroBioStand).

## Notes

### Competing Interest Statement

The authors have declared no competing interest.

